# AnchorR: A QuPath and R interface for collaborative exploration of spatial transcriptomics and histology

**DOI:** 10.64898/2026.08.05.742985

**Authors:** Caitlin A Morris, William C Bastian, Yan Cui, Zoya Kurago, Eugene F Douglass

## Abstract

Single-cell spatial transcriptomics can connect molecular cell states with tissue morphology, but this promise depends on accurate registration to histopathology. In serial sections, however, tissue borders often differ because of sectioning artifacts, staining variability, and field-of-view acquisition, limiting conventional area-based registration. We developed AnchorR, an expert-guided workflow for coarse-grained alignment of hematoxylin and eosin (H&E) images with CosMx Spatial Molecular Imaging data. Bioinformaticians first define and color-code cell types in Seurat, and pathologists then identify corresponding internal landmarks using QuPath overlays. AnchorR combines these paired landmarks to estimate affine transformations, quantify residual error, and support visual quality control and anchor refinement.

Using six oral pre-cancerous tissue sections, we identified 60 cross-modal landmarks. Fitting each section independently reduced mean landmark error from 121.5 µm with a single whole-slide transformation to 14.6 µm. Cross-validation further showed that increasing the number of anchors improved robustness, with nine-anchor fits achieving approximately 20 µm error, or about one cell diameter.

AnchorR is designed to complement automated computer-vision methods by providing reliable tissue-level alignment when border mismatch makes global registration difficult. By creating a shared workspace for pathologists and bioinformaticians, it operationalizes an expert-in-the-loop approach and makes feature-based multimodal registration accessible without specialized computer-vision expertise or high-performance computing.

## Introduction

### Single-cell spatial omics

is an emerging technology that could unite immunology, genomics, and histopathology by linking molecular and immune states directly to tissue morphology.^1, 2^ This integration could reveal new biology and improve the diagnosis of histologically ambiguous lesions,^3^ but different processing requirements often mean that each modality must be applied to adjacent serial sections.^4^

### Serial sectioning

introduces deformation at nearly every processing step: fixation and dehydration shrink and harden tissue; embedding determines section orientation and can promote curling or bending; microtomy can compress, shear, tear, or fold sections; and flotation, mounting, drying, and staining can further stretch, wrinkle, or locally distort the tissue.^5, 6^ Consequently, multimodal analysis across serial sections depends critically on image registration that can correct not only differences in position, orientation, and scale, but also physical deformations such as tears and shear.^7^

### Image registration

is the process of geometrically overlaying two or more images of the same scene, acquired at different times, viewpoints, or with different sensors, so that their information can be analyzed jointly.^8^ Many foundational concepts were established in a landmark 1992 survey,^9^ after which registration was increasingly applied across fields including satellite imaging and medicine. ^8^ Methods broadly fall into two classes: *<u>area-based approaches</u>*, which align images from their overall intensity or geometric patterns without identifying specific structures, and *<u>feature-based approaches</u>*, which match landmarks such as corners, centroids, intersections, or endpoints and require expert annotation.^7, 10, 11^ Historically, radiologic and histologic registration has relied largely on area-based methods because medical images often lack abundant, consistently identifiable landmarks. ^8^

### Current medical pipelines

simplify whole-slide images so that global registration is driven mainly by gross tissue shape rather than internal histologic structure. ^7, 10, 11^ Images are typically converted to grayscale, down-sampled, separated from background, and coarsely aligned by translation, rotation, scaling, or affine transformation, making the outer tissue border the dominant signal.^12, 13^ Recent benchmarks comparing hematoxylin and eosin (H&E) with immunohistochemistry (IHC) have produced two consistent conclusions: *<u>border mismatch is the leading cause of registration failure (Fig 1A)</u>*, and feature-based methods perform best when such mismatch occurs (Fig 1C).^7, 10, 11^ Border mismatch can arise from tears, inconsistent staining, or tissue cropping^5, 6^ and is especially pronounced in single-cell spatial transcriptomic platforms such as CosMx Spatial Molecular Imaging (SMI), where field-of-view (FOV) acquisition creates artificial boundaries and immunofluorescence (IF) membrane staining delineates tissue differently from H&E (Fig 1A).^1^ Feature-based methods avoid this dependence on outer contours by matching internal structures, but reliable landmark selection often requires expert histopathologic and bioinformatic annotation (Fig 1B).^8^

**Figure 1.**
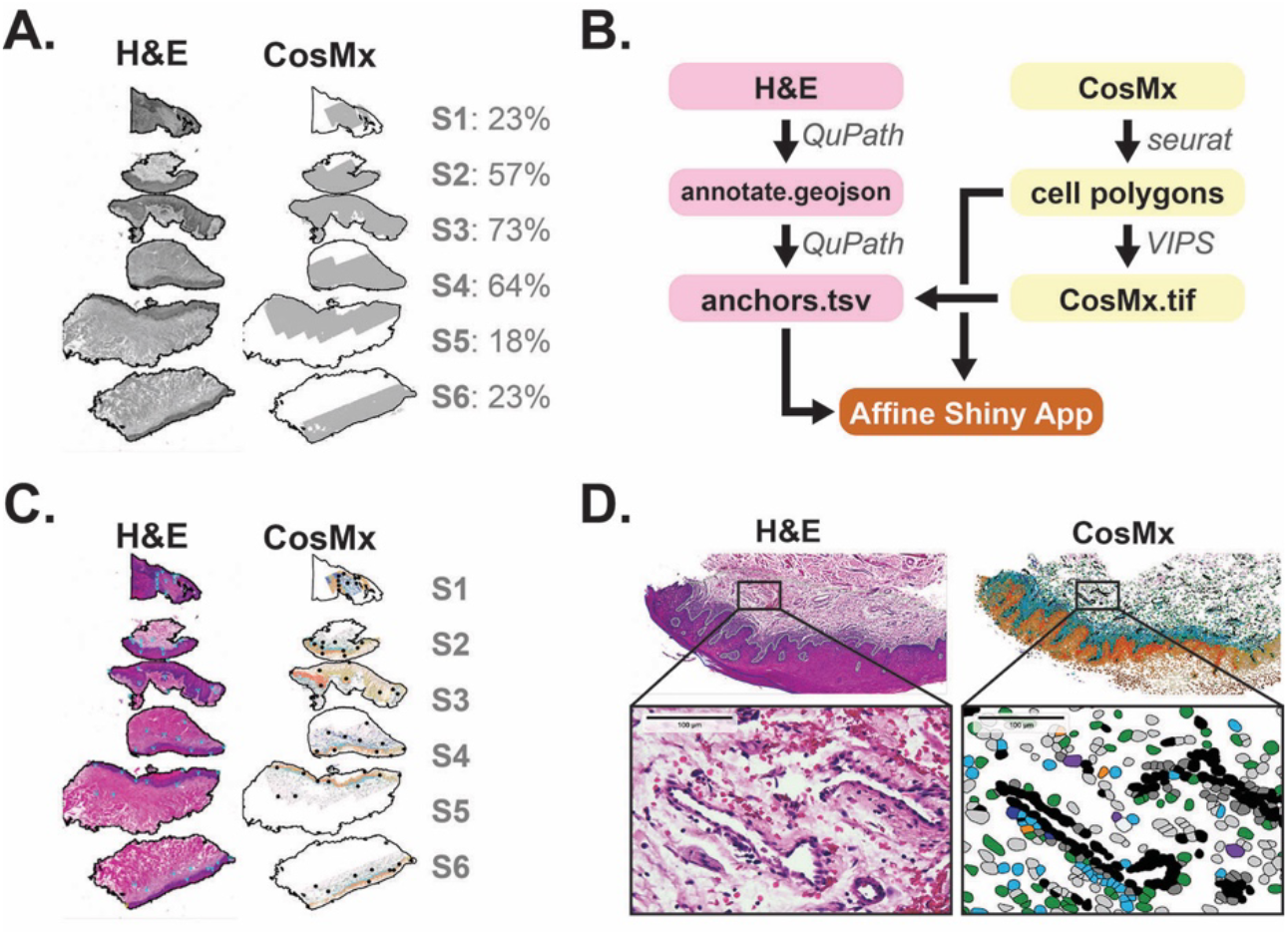
Expert-guided registration of serial H&E and CosMx spatial transcriptomic sections using AnchorR. **(A)** Grayscale CosMx cell polygons and corresponding H&E sections from a WSI containing six oral pre-cancerous biopsies. H&E borders were defined by thresholding in QuPath, and CosMx polygons were transformed using the AnchorR-derived affine registration. Border overlap was calculated as the fraction of the H&E boundary with a CosMx polygon within 200 µm. **(B)** AnchorR workflow. Pathologists annotate epithelial-stromal boundaries in QuPath and export a GeoJSON file, while bioinformaticians assign and color-code cell types in Seurat to generate a cell-polygon file. The polygons are converted to an image and overlaid with H&E, enabling identification of recurrent cross-modal structures and export of H&E landmark coordinates as an anchors.tsv file. AnchorR then determines the corresponding CosMx coordinates and estimates the affine transform. **(C)** Workflow output showing 10 paired landmarks per tissue section. **(D)** Representative endothelial landmark, demonstrating that vascular structures provide recurrent features beyond epithelial-stromal boundaries.

Overall, in our experience, the central challenge in registering histopathology with spatial transcriptomics is border mismatch.^10, 11^ Figure 1A shows a whole-slide image (WSI) containing six oral pre-cancer biopsies, with substantial variation in overlap between H&E tissue boundaries and those defined by CosMx field-of-view (FOV) acquisition and membrane staining. This variability makes high-fidelity global alignment difficult, and often intractable, for area-based methods.^13, 14^ Feature-based approaches are more robust but require landmarks defined jointly by pathologists and messenger RNA (mRNA) bioinformaticians, two communities that rarely have computer-vision expertise.^4, 15^

### We therefore developed AnchorR

a collaborative pipeline and interactive R-based tool that connects these experts through software already familiar to them: QuPath for histopathology^12^ and R for spatial transcriptomic analysis (Fig 1B). By enabling direct communication between pathologists and bioinformaticians, AnchorR supports the identification of biologically corresponding landmarks visible in both modalities. Importantly, it requires neither specialized computer-vision expertise nor computationally intensive infrastructure,^4, 15-20^ democratizing cross-modal analysis while answering recent calls for “experts in the loop.”^7^

## Results

### Seurat Workflow: defining cell types

Histopathologic landmarks are well established and include tissue-border corners, endpoints, and centroids of structures such as ducts, vessels, and villi. ^7, 10, 11^ Defining analogous landmarks in spatial transcriptomics is less straightforward because single-cell spatial transcriptomics primarily provides cell boundaries with limited intracellular structural information.^1^

The cell is therefore the fundamental spatial unit,^1^ and many current registration approaches rely on unsupervised gene-expression clustering as the primary source of cell-level information.^4, 15-20^ This is often sufficient to distinguish epithelial and stromal compartments and has supported registration of serial sections from single-sample whole-slide images (WSIs).^4, 16, 19^ In multi-sample WSIs, however, batch effects can fragment epithelial and stromal populations into sample-specific subclusters (Figure S1).^21^ Additional supervision is therefore needed to identify sample-level batches and assign biologically interpretable cell types rather than relying on transcriptional clusters alone. These steps require bioinformatic interpretation beyond computer vision alone.

As detailed in the Supporting Information tutorial and accompanying R code, we developed an adapted Seurat workflow for single-cell spatial transcriptomic data(Fig 1B, “Seurat”).^22^ Key steps include standard quality control, visual identification of tissue-section batches, Harmony batch correction, clustering, and supervised cell-type annotation. The final output is a cell-polygon file color-coded by cell type, which is used to identify corresponding structures in hematoxylin and eosin (H&E) images and define CosMx anchors(Fig 1D). We provide complete code, example data, and a step-by-step tutorial because Seurat-based methods for single-cell spatial transcriptomics are evolving rapidly and are often difficult to reconstruct from the literature or outdated online tutorials.

### Defining multi-modal landmarks

With the color-coded cell polygons defined, the next step is to convert them into a format that can be viewed in standard digital pathology software such as QuPath.^12^ Because whole-slide imaging platforms operate on tiled image data, the cell polygons are first rendered as a PNG image and then converted into a tiled TIFF compatible with QuPath overlays. This conversion is performed using VIPS, an open-source image-processing library designed for efficient handling of large images (Figure 1B, “VIPS”).^23^ The resulting image preserves the spatial distribution and cell-type color coding of the CosMx data while allowing direct overlay with the corresponding hematoxylin and eosin image.

Using the QuPath Align extension, the H&E and CosMx images are imported into a shared project and their pixel scales approximately matched (Fig 1B “QuPath”). For each tissue section, the CosMx overlay is displayed at approximately 50% opacity and manually rotated, translated, and, when necessary, rescaled to obtain a coarse alignment based on the tissue border and one or two internal structures. The pathologist then defines 10 landmarks distributed across the section, prioritizing distinctive structures such as ducts, vessels, and epithelial islands, followed by corners or tissue tips when needed(Fig 1C). Each landmark is verified by toggling between the H&E and CosMx views to confirm that it is visible in both modalities; endothelial structures provided particularly reproducible cross-modal landmarks (Fig 1D). The final H&E landmark coordinates are exported as a tab-separated values (TSV) file for AnchorR, with screenshots retained for quality control.

### AnchorR affine registration tool

R is widely used for bioinformatic and statistical analysis but is not naturally suited to visual landmark selection because it lacks a native graphical user interface (GUI) for interactive image overlays. We therefore developed AnchorR, an R Shiny application that combines CosMx anchors identified from the cell-type map with QuPath point exports and H&E tissue borders to support visually guided landmark placement, affine registration, and quality control (Figure 1B). The app enables users to define and refine corresponding landmarks without specialized computer-vision software or programming expertise.

Tab 1 supports CosMx anchor placement using a two-panel interface: a global cell-type map is used to navigate the slide, while a higher-resolution local view enables precise landmark placement within a selected field-of-view neighborhood. Tab 2 matches these CosMx coordinates to QuPath-derived H&E landmarks by identifier, estimates the affine transformation, and reports its rotation, scale, stretch, shear, translation, and fitting error. Registration quality is displayed through predicted-versus-observed anchor positions, per-anchor residuals, and transformed H&E tissue borders overlaid on the CosMx map. Individual landmarks can be excluded and the model refitted, allowing rapid visual identification and correction of poorly defined anchors before exporting the final affine matrix.

A detailed tutorial is provided in the Supporting Information, and the source code is available on GitHub. AnchorR can be applied to any dataset that provides standard cell_meta.csv and cell_polygons.csv inputs. At minimum, the metadata file must contain “cell” and “color” columns, linking cell identifiers to display colors, while the polygon file must contain “cell”, “x_global_px”, and “y_global_px” columns defining each cell boundary in global image coordinates.

### WSI versus individual-section registration

Following the Supporting Information tutorial, we identified 60 landmarks conserved across the CosMx and H&E modalities, with 10 landmarks placed in each of six tissue sections (Fig 1C). This balanced representation prevented larger sections from dominating model fitting and provided additional landmarks for cross-validation and sensitivity analyses. We first tested whether a single affine transformation fitted across the entire whole-slide image (WSI) or separate transformations fitted within each tissue section produced better registration (Fig 2). Section-specific fitting substantially outperformed whole-slide registration. The mean per-anchor error decreased from 121.5 µm with a single WSI-level affine to 14.6 µm with independent section-level affines, representing an approximately eightfold improvement and nearly an order-of-magnitude reduction in error (Fig 2B). This result is consistent with recent studies that regional registration on WSI can out-perform whole-slide registration.^24^ Accordingly, all subsequent registration, cross-validation, and error analyses were performed independently within each tissue section.

**Figure 2.**
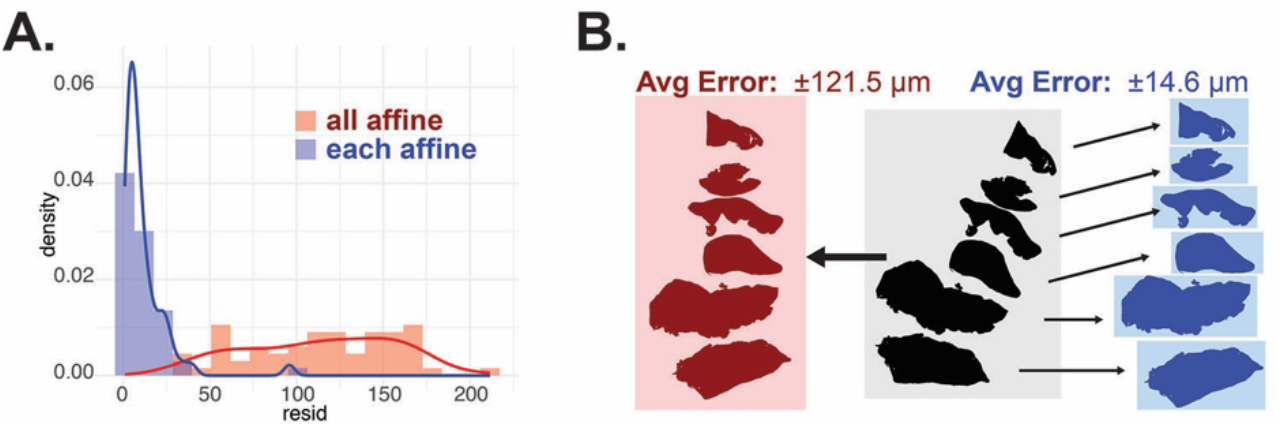
Section-specific affine registration improves alignment compared with whole-slide fitting. **(A)** Distribution of per-landmark registration residuals obtained using a single affine transformation fitted across all six tissue sections (“all affine,” red) or separate affine transformations fitted independently within each section (“each affine,” blue). Section-specific fitting produced substantially lower residuals. **(B)** Schematic comparison of whole-slide image (WSI)-level and section-level registration. A single global affine transformation yielded a mean registration error of 121.5 µm, whereas independent section-specific affine transformations reduced the mean error to 14.6 µm. These results indicate that processing-induced changes in orientation, scale, and deformation vary across tissue sections and are not adequately captured by a single WSI-level transform.

### Sensitivity analysis of registration accuracy

We next quantified registration error using a test-point error cross-validation strategy^8^ in which affine transformations were fitted using three to nine landmarks and evaluated on the held-out points. This analysis was performed across all possible landmark combinations, with the mean test root-mean-square error (RMSE) and interquartile range summarized in Figure 3A. Assuming that corresponding landmarks were correctly identified across modalities, the results suggest that individual landmark-placement error was approximately 20-60 µm, while overall fitting error decreased as additional anchors averaged out uncertainty in individual coordinates. With nine anchors, test RMSE fell to approximately 20 µm, or roughly one cell diameter, which is sufficient for tissue-level analyses.

**Figure 3.**
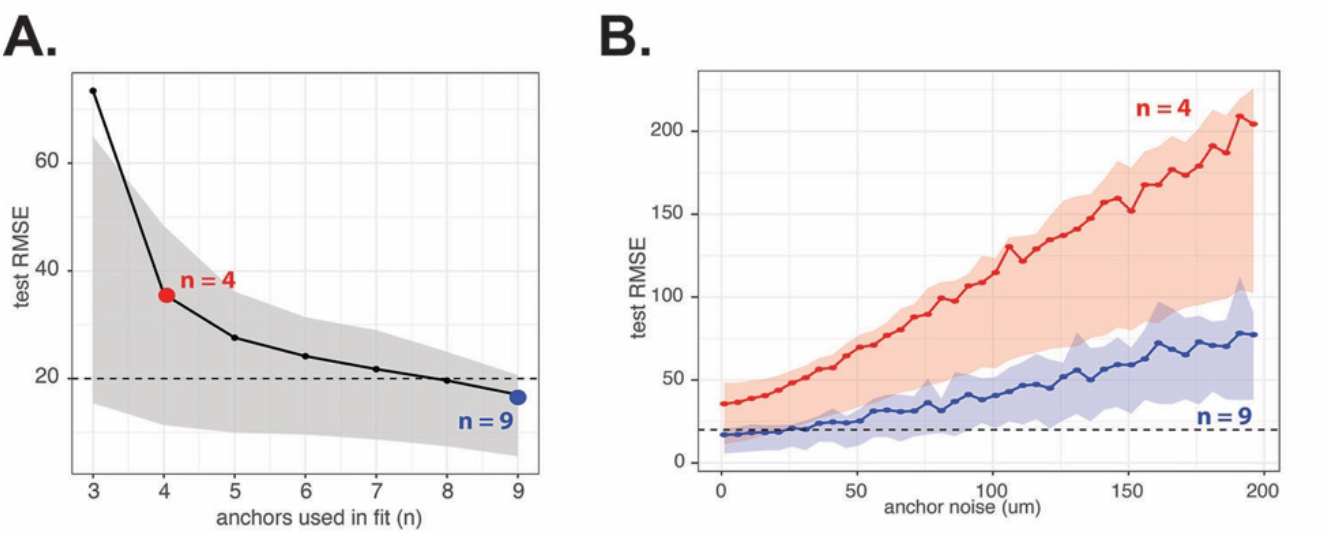
Cross-validation and sensitivity analysis of affine registration accuracy. **(A)** Test-point error was estimated by fitting affine transformations with three to nine anchors and evaluating root-mean-square error (RMSE) on the held-out landmarks. All possible anchor combinations were assessed, and the mean test RMSE is shown with the interquartile range. Registration error decreased as additional anchors were included, reaching approximately 20 µm with nine anchors. **(B)** Robustness to landmark-placement uncertainty was evaluated by repeating the test-point procedure after adding increasing amounts of synthetic noise to the anchor coordinates. Fits using nine anchors (blue) consistently produced lower test RMSE than fits using four anchors (red), including at high noise levels. Shaded regions indicate the interquartile range, and the dashed horizontal line marks a 20 µm reference error, approximately one cell diameter.

To assess robustness to increasing annotation uncertainty, we repeated the procedure after adding synthetic noise to each landmark (Fig 3B). Fits using nine anchors consistently outperformed those using four anchors and remained substantially more accurate even at 200 µm of simulated noise, reducing error by approximately threefold. This improvement is consistent with least-squares theory: because a two-dimensional affine transformation estimates three coefficients per coordinate, increasing the number of well-distributed anchors reduces parameter-estimation variance approximately in proportion to 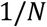, and registration error approximately in proportion to 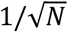. The exact reduction depends on anchor geometry and target location.^9^

## Discussion

Recent benchmarking studies have highlighted a persistent limitation of multimodal histology registration: area-based methods perform well when tissue borders and staining patterns are preserved, but often fail when serial sections contain tears, cropping, stain-dependent boundaries, or other processing artifacts.^7, 10, 11^ Feature-based approaches are more robust because they align internal structures rather than relying on global image similarity, yet their broader use has been limited by the need for biologically meaningful landmarks.^8^ In histopathology-spatial transcriptomics registration, these landmarks cannot be defined reliably by computer vision alone because they must be recognizable both morphologically (= pathologist expertise) and molecularly (= bioinformatics expertise).

AnchorR addresses this gap by creating a shared workspace in which pathologists and bioinformaticians jointly define cross-modal landmarks using familiar tools (Fig 1B). The workflow is intended to complement, rather than replace, automated computer-vision registration. It provides coarse-grained affine alignment suitable for tissue compartments, regional patterns, and multicellular structures, but not pixel-level or single-cell correspondence. Its primary advance is therefore not a new high-resolution registration algorithm, but an accessible framework for incorporating domain expertise into the most failure-prone stage of multimodal alignment. By formalizing collaboration between histopathology and bioinformatics, AnchorR fits naturally within an expert-in-the-loop framework and broadens access to feature-based registration for teams without specialized computer-vision expertise.

The final tutorial step transforms the complete cell-polygon dataset into hematoxylin and eosin (H&E) coordinate space, enabling downstream bioinformatic analyses to be performed in direct proximity to the pathologist-defined histologic reference. This supports regional comparisons between molecular cell states and tissue morphology while preserving the original single-cell spatial data structure. We also provide code and instructions for an OpenSeadragon interface that enables direct overlay and comparison of H&E and spatial transcriptomic layers through web-based and mobile-compatible viewers, further supporting collaborative review across modalities.

## Conclusion

AnchorR provides an accessible expert-guided workflow for coarse-grained registration of H&E and single-cell spatial transcriptomic data across serial sections. By combining pathologist-defined morphology with bioinformatic cell-type annotation, the approach enables robust internal landmark selection when border mismatch limits conventional area-based methods. Section-specific affine fitting and multi-anchor placement support tissue-level alignment while preserving a clear distinction from pixel-level or single-cell registration. The resulting transformation places cell polygons directly into H&E space, allowing downstream spatial analyses to remain anchored to histopathologic ground truth. Together with QuPath, R, and web-based visualization tools, AnchorR establishes a practical expert-in-the-loop framework that strengthens collaboration between pathology and bioinformatics and broadens access to multimodal tissue analysis.

## Supporting information

Detailed Method Protocol

## Acknowledgements

This study is supported in part by NIH/NIDCR 1R01DE032178 to YC and the Georgia Cancer Center Paceline funding mechanism MCGFD01068 to YC and ZK. We acknowledge the support and contribution of the Integrated Genomics Core Shared Resources at the Georgia Cancer Center, Augusta University (RRID: SCR_026483) for this study

## Lead Contact

Further information and requests for resources should be directed to and will be fulfilled by the Lead Contact, Eugene Douglass.

## Materials Availability

This study did not generate new unique reagents.

## Data and Code Availability

The analysis, prototype code and datasets are available through a GitHub repository: http://www.github.com/fingolfn/AnchorR Raw data is available through Figshare. both will be made publically available upon final publication.

## Author Contributions

ED, ZK, and YC conceived the study, developed the analytical framework. ZK performed all histopathological annotation and provided clinical guidance. CM and ED performed all computational analyses. CM developed the first OpenSeaDragon prototypes. WB developed the CosMx data processing pipeline. ED developed the registration pipeline and AnchorR tool. All authors contributed manuscript editing and approved the final manuscript.

## Methods

### SAMPLES: Tissue specimens and multimodal imaging

#### Sample acquisition and preparation

The study was approved by the AU IRB (Protocol 964456) and funded by the internal 2024 Paceline grant. Formalin-fixed paraffin-embedded archival specimens were selected based upon diagnosis, quality, and quantity of available material from the Oral and Maxillofacial Pathology Laboratory at Augusta University Dental College of Georgia. Selected samples were assembled into a block meeting the 2 x 1.5 cm dimensions required for CosMx processing. Seven serial 4.5 um sections were mounted on VWR Superfrost Plus Slides; slide #4 was stained by H&E, while all other sections were air-dried.

#### COSMX: slide preparation and spatial data acquisition

The six unstained serial sections were processed for CosMx SMI following the manual FFPE RNA slide-preparation protocol (Bruker Spatial Biology, MAN-10184-07), with all six sections mounted inside the 20 × 15 mm imageable scan area of a single slide. Slides were baked overnight at 60 °C, deparaffinized in xylene and ethanol, and subjected to target retrieval for 15 min at 100 °C, followed by Proteinase K digestion (3 µg/mL, 30 min at 40 °C). Fluorescent fiducials were applied at 0.001% in 2X SSC-T and must be present on bare glass within the scan area, which is why sections were spaced to leave uncovered glass between them. Tissue was post-fixed in 10% neutral-buffered formalin, blocked with 100 mM NHS-acetate, and hybridized overnight at 37 °C with the CosMx Human 6K Discovery RNA panel. After two 25-min stringent washes (1:1 formamide/4X SSC, 37 °C), sections were stained with DAPI and the CosMx Human Universal Cell Segmentation markers (CD298/B2M, PanCK, CD45) [CONFIRM], assembled into flow cells, and imaged on the CosMx SMI instrument. Barcodes were read out by cyclic in situ hybridization across user-selected fields of view, and morphology images in the DAPI and segmentation-marker channels were used for on-instrument cell segmentation. Fiducials provide focus and cycle-to-cycle image registration within a run; they do not relate the CosMx frame to any external image, which is the gap AnchorR addresses. Raw data were processed in the AtoMx Spatial Informatics Platform, from which the Seurat object and cell-boundary flat files described below were exported. All CosMx cell coordinates and polygon vertices are reported in the global pixel coordinate system of the slide (0.2501 µm/pixel), and it is this frame that the affine transformation maps into H&E space.

### SEURAT: CosMx preprocessing and cell-type annotation

#### Quality Control

A Seurat object generated from the CosMx dataset using the AtoMx Spatial Informatics Platform was imported into R. The object contained 195,069 cells and 6,175 measured genes. Individual tissue sections were identified from their spatial positions on the slide using the x_slide_mm and y_slide_mm coordinates, and cells were assigned to one of six sections (S1–S6). Cells were retained only when they passed both cell-level quality control (qcCellsPassed = TRUE) and field-of-view quality control (qcFlagsFOV = ''Pass”), resulting in 191,704 cells for downstream analysis.

#### Batch Correction and clustering

Gene-expression data were normalized, variable features were identified, and expression values were scaled using Seurat with default settings. Principal-component analysis was performed using 30 components, followed by Harmony correction across the six tissue sections using sample_slide as the batch variable. The first 30 Harmony dimensions were used for UMAP visualization, nearest-neighbor graph construction, and graph-based clustering at a resolution of 0.5, producing 21 transcriptional clusters. Positive cluster markers were identified using FindAllMarkers, requiring expression in at least 25% of cells and an average log2 fold-change greater than 0.25. The 30 markers with the highest average log2 fold-change in each cluster were used for annotation.

#### Cell-type definitions

Clusters were assigned to biologically interpretable cell types through supervised evaluation of canonical marker genes and comparison with annotations from a previously characterized sample. The resulting labels included basal and squamous epithelial states, keratinized epithelial cells, T cells, B cells, macrophages, dendritic cells, mast cells, fibroblasts, endothelial cells, pericytes, and muscle cells. Ambiguous clusters were retained as low-confidence or unknown populations rather than excluded. Final cell-type assignments were stored in the Seurat metadata and associated with a consistent color palette for spatial visualization and generation of the cell-polygon registration image.

### VIPS: Cell-polygons and image generation

#### Pre-registration: preparation of QuPath and AnchorR inputs

Cell-boundary polygons were imported from the CosMx flat-file output and filtered to retain only cells present in the quality-controlled, cell-type-annotated Seurat object. Cell identifiers were used to transfer the final cell-type assignments and associated display colors from the Seurat metadata to each polygon (Fig 1C-D). Two input files were then generated for AnchorR: cell_polygons.csv, containing the polygon vertices in CosMx global pixel coordinates, and cell_meta.csv, containing the cell identifier, field of view, final cell-type assignment, and display color.

For visualization in QuPath,^12^ polygons were rendered in their original CosMx coordinate system using cell type as the fill color and cell identifier to group vertices into individual polygons. The resulting image was exported as a high-resolution PNG. VIPS, an open-source image-processing library optimized for efficient handling of large images,^23^ was then used to flatten the image onto a white background and convert it into a tiled, pyramidal TIFF using 256 × 256-pixel tiles and LZW compression. The pyramidal TIFF was imported into QuPath and, when required, exported as an OME-TIFF at the original pixel resolution for use as an overlay during landmark identification.

#### Post-registration: cell polygons and images in H&E space

Following affine registration, each cell polygon was assigned to its corresponding tissue section using the section labels stored in the Seurat metadata. Separate CosMx-to-H&E affine matrices were loaded for each of the six tissue sections. For every polygon vertex, the CosMx global coordinates were expressed in homogeneous form as [x CosMx ,y CosMx ,1] and multiplied by the corresponding section-specific affine matrix. The transformed coordinates were stored as x_he_px and y_he_px, thereby placing the complete cell-polygon dataset directly into the H&E pixel-coordinate system. The resulting polygon table, including the original coordinates, section labels, cell types, and transformed H&E coordinates, was saved as an R data object for downstream spatial analyses.

The transformed polygons were also rendered as a PNG using the exact width and height of the H&E whole-slide image as the plotting coordinate limits. The vertical axis was reversed to reproduce the top-left origin used by QuPath, and plot expansion and margins were removed to preserve correspondence between image and data coordinates. Because direct full-resolution rendering exceeded available memory, the raster image was exported at one-third of the H&E linear resolution while retaining the full H&E coordinate extent. The resulting PNG could subsequently be resized to the native H&E dimensions or converted with VIPS into a tiled, pyramidal TIFF for efficient overlay and visualization.

### QUPATH: H&E border and landmark annotation in QuPath

#### H&E tissue boundaries

were identified in QuPath using a pixel-intensity thresholding procedure adapted from the QuPath tissue-detection workflow. A thresholder was applied across the entire image at 1.10 µm/pixel using the Eosin channel, Gaussian prefiltering with a smoothing sigma of 5, and an intensity threshold of 0.01. Pixels above the threshold were classified as tissue (“Tissue”), whereas pixels below the threshold were left unclassified. The thresholded regions were converted into annotation objects using a minimum object area of 200,000 µm^2^ and a minimum enclosed-hole area of 10,000 µm^2^. Disconnected regions were split into separate objects, and existing threshold-derived annotations were replaced. The resulting annotations were visually inspected to confirm that they followed the outer tissue contours and were exported from QuPath as GeoJSON polygons for use in AnchorR registration and border-overlap analyses.

#### Multi-modal Landmark annotation

H&E and CosMx cell-type images were imported into a shared QuPath project and approximately scale-matched using the QuPath Align extension. The H&E image was used as the fixed reference, while the CosMx layer was displayed at approximately 50% opacity and manually rotated, translated, and, when necessary, rescaled for each tissue section. Coarse alignment was guided by the tissue contour and one or two large internal structures.

Ten corresponding landmarks were then placed across each section and labeled L1-L10. Landmarks were distributed across both ends and the center of the tissue, prioritizing distinctive ducts, vessels, and epithelial islands, followed by corners or tissue tips when necessary. Candidate landmarks were verified by toggling between the H&E and CosMx layers to confirm that the same structure was visible in both modalities. Landmark coordinates were exported from QuPath as a TSV file, and paired H&E and CosMx screenshots were retained for quality control. A full detailed procedure for this manual process is provided in the supporting information.

### AnchorR: Application and software implementation

AnchorR was implemented as a two-tab R Shiny application using ggplot2, data.table, sf, and related R packages. *<u>Tab 1: CosMx anchor definition</u>* loads the quality-controlled cell_polygons.csv and cell_meta.csv files and displays the color-coded cell polygons in their original CosMx global pixel coordinates. A cached whole-slide overview supports navigation, while a higher-resolution panel displays the selected field of view and its surrounding 3 × 3 field-of-view neighborhood. Users place sequentially labeled landmarks by clicking the detailed cell-type map and can review, undo, clear, or export the resulting CosMx coordinates as a CSV file containing landmark identifier, x-coordinate, and y-coordinate.

*<u>Tab 2: affine fitting and quality control</u>* imports the corresponding H&E landmarks exported from QuPath as a TSV file and optionally imports H&E tissue-border polygons as GeoJSON. CosMx landmarks can be passed directly from Tab 1 or uploaded separately. Landmarks are paired by identifier, and a two-dimensional affine transformation is estimated by least-squares regression using at least three matched landmark pairs. The transformation can be fitted in either the H&E-to-CosMx or CosMx-to-H&E direction and is summarized by its translation, rotation, scale, shear, determinant, root-mean-square error, and maximum residual. Registration fidelity is evaluated using predicted-versus-observed landmark positions, two-dimensional residual plots, and per-landmark errors. Individual landmarks can be excluded interactively and the model automatically refitted. For visual quality control, the fitted transformation is applied to the H&E tissue-border polygons and the transformed borders are overlaid directly on the CosMx cell-type map. The final affine matrix, transformed GeoJSON borders, and overlay image can then be exported for downstream analysis.

### Registration performance evaluation

Registration performance was evaluated using 60 paired landmarks, with 10 landmarks defined in each of six tissue sections. Two affine-fitting strategies were compared: a single whole-slide transformation fitted to all 60 landmark pairs and six section-specific transformations fitted independently to the 10 landmarks within each section. Affine matrices mapping CosMx coordinates to H&E coordinates were estimated by ordinary least squares. For each landmark, registration error was calculated as the Euclidean distance between its observed H&E position and the position predicted from its CosMx coordinates. Residual distributions and root-mean-square error (RMSE) were compared between the whole-slide and section-specific models.

### Cross-validation and sensitivity analysis

Registration accuracy was assessed by exhaustive landmark-dropout cross-validation within each tissue section. Affine transformations were fitted using every possible combination of three to nine of the 10 available landmarks, and performance was evaluated on the landmarks excluded from each fit. Training and held-out RMSE were calculated as the root mean squared Euclidean distance between predicted and observed coordinates. Results were pooled across sections and combinations and summarized by the mean and interquartile range for each number of fitting landmarks.

Sensitivity to landmark-placement uncertainty was evaluated by independently adding normally distributed noise to the x- and y-coordinates of the CosMx landmarks used for model fitting, while leaving held-out landmarks unchanged. Noise levels ranged from 1 to 200 µm in 5-µm increments, and the complete cross-validation procedure was repeated at each level using a fixed random seed. Robustness was summarized by comparing the mean and interquartile range of held-out RMSE for transformations fitted with four versus nine landmarks.

