## Supplementary material for "AnchorR: A QuPath and R interface for collaborative exploration of spatial transcriptomics and histology": Detailed Method Protocol

### Seurat: CosMx Pre-processing and cell-type annotation

```
####################################################################################################################################
####################################################################################################################################
##     ########      ########      ########     ########             #########           ##       ############      ##
##     ##     ##     ##     ##     ##           ##     ##            ##      ##         ####           ##          ####
##     ##     ##     ##    ##      ##           ##     ##            ##       ##       ##  ##          ##         ##  ##
##     ########      #######       ########     ########             ##        ##     ##    ##         ##        ##    ##
##     ##            ##    ##      ##           ##                   ##       ##     ##########        ##       ##########
##     ##            ##     ##     ##           ##                   ##      ##     ##        ##       ##      ##        ##
##     ##            ##      ##    ########     ##                   #########     ##          ##      ##     ##          ##
####################################################################################################################################
####################################################################################################################################
##       ########    ########   ##    ##   #######         ##     ##########
##      ##           ##         ##    ##   ##    ##       ####          ##
##      ##           ##         ##    ##   ##    ##      ##   ##        ##
##       #######     #######    ##    ##   #######      ##     ##       ##
##             ##    ##         ##    ##   ##    ##    ##########       ##
##             ##    ##         ##    ##   ##     ##   ##        ##     ##
##       #######     #######     ######    ##      ##  ##        ##     ##
####################################################################################################################################
####################################################################################################################################
#GOAL:  full processing through cell-types (to define internal landmarks)
```

This procedure processes raw CosMx expression data into a quality-controlled and annotated Seurat object. The input is an AtoMx-derived Seurat object containing gene-expression data, spatial coordinates, and QC metadata. Cells are assigned to individual tissue samples, filtered using cell- and FOV-level QC criteria, normalized, batch-corrected with Harmony, and clustered based on gene-expression profiles. Cluster marker genes are then used to assign biological cell types, which are validated in both UMAP and tissue space. The final output is a processed Seurat object containing corrected embeddings, cluster assignments, and spatially resolved cell-type annotations for downstream registration and spatial analysis.

#### Quality Control of Raw Data

1. **Import the CosMx Seurat object and inspect existing results.**
   1.1. Load the Seurat object exported from AtoMx and review its dimensions, metadata columns, and previously generated analyses.
   1.2. Confirm the number of cells and genes, summarize cell- and FOV-level quality-control results, and visualize the existing AtoMx cell clusters and spatial neighborhood assignments in both UMAP and slide-coordinate space.
      1.2.1. Look for clear evidence of batch effects:
      - Many discrete clusters (as seen below in sample data). CosMx data generally doesn't appear this cluster-y in the absence of batch effects (usually only two major clusters with smearing between sub clusters)

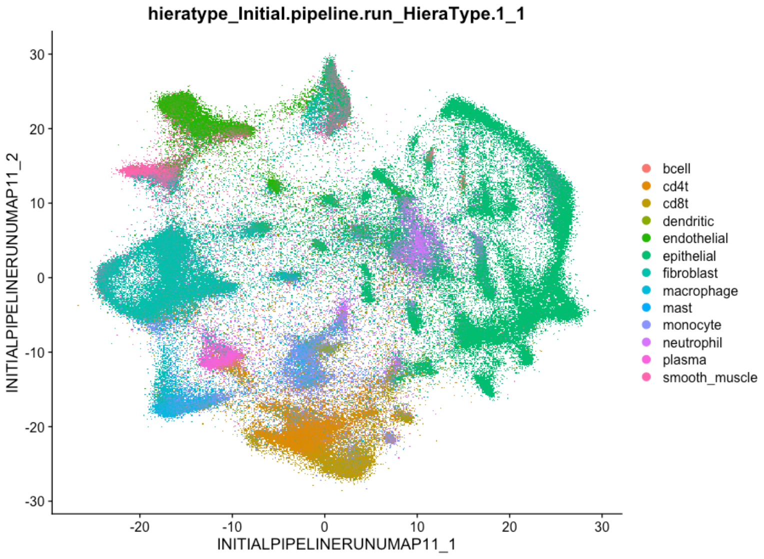

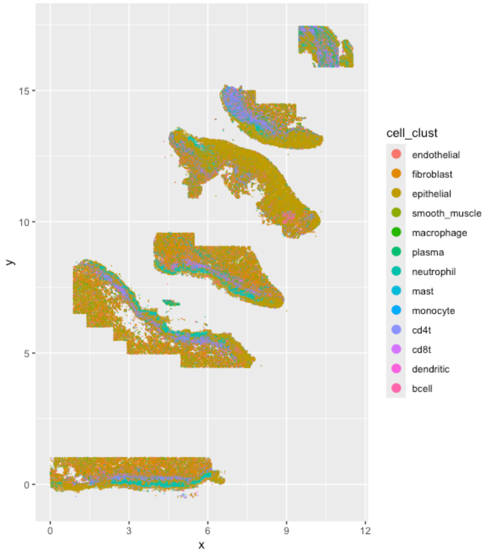

2. **Define the individual tissue samples on the CosMx slide.**

2.1. Plot cell coordinates using x_slide_mm and y_slide_mm to identify the spatial boundaries separating the six tissue sections.

2.2. Plot histogram along major sample discriminating axis (here y) to identify rough horizontal thresholds:

- hist(nano.obj@meta.data$y_slide_mm,breaks=100)

2.3. Define one horizontal and four angled boundary lines, using abline:

2.3.1. a = intercept or rough horizontal gap

2.3.2. b = slope

- abline(h=c(3),lty=2,lwd=2,col="red")
- abline(a=19,b=(-0.4),col="blue",lty=2,lwd=2)#angle 1

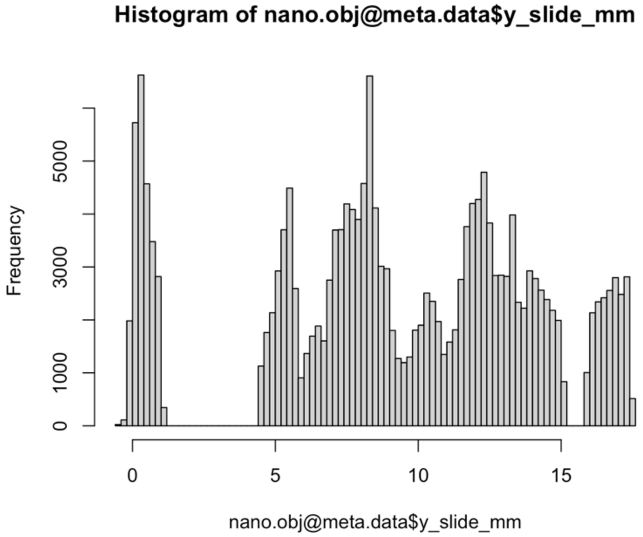

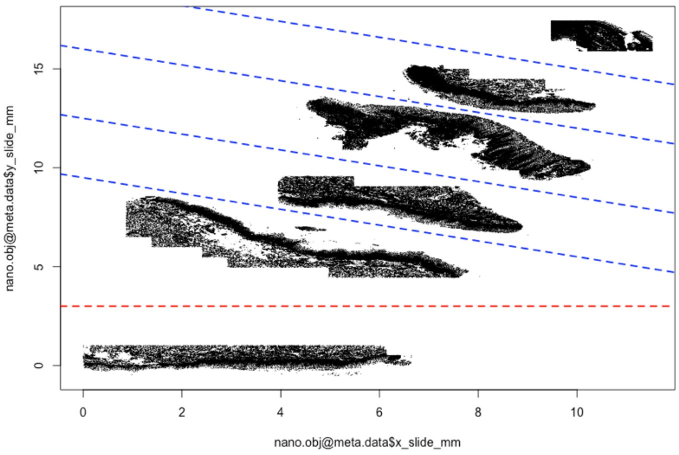

2.4. Assign each cell to samples S1–S6 based on its position relative to these thresholds. Visually inspect the resulting sample labels in slide space to confirm that the tissue sections were separated correctly.

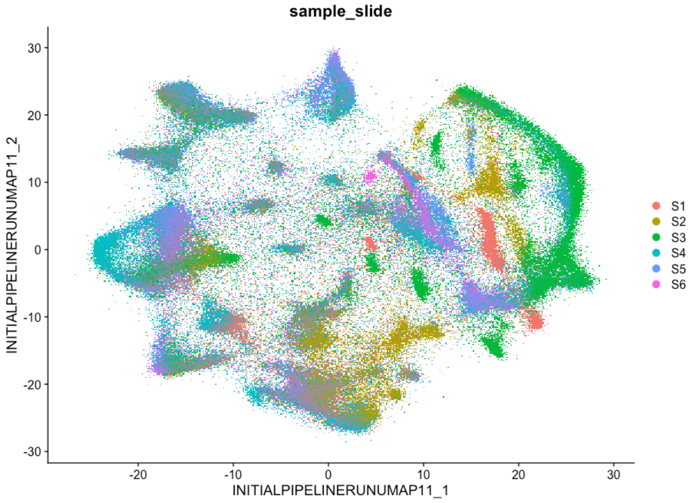

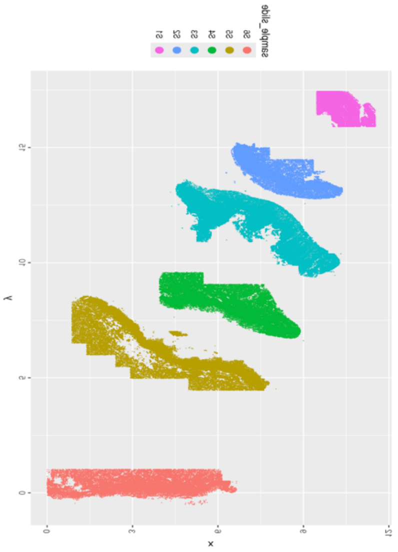

3. **Evaluate and apply cell- and FOV-level quality control.**

3.1. Examine the distributions of qcCellsPassed and qcFlagsFOV in both UMAP and physical slide space. Calculate a log-transformed transcript-detection measure from nFeature_RNA to identify regions with unusually low expression depth. Remove the image slot before subsetting, then retain only cells that passed cell-level QC and originated from FOVs labeled as passing. Record the number of cells retained after filtering.

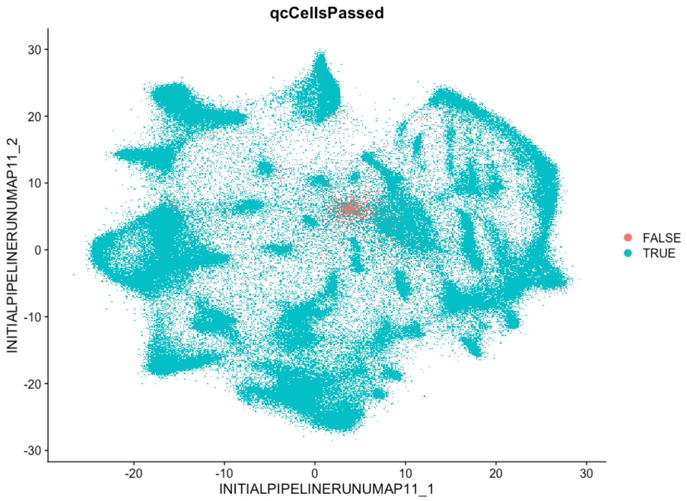

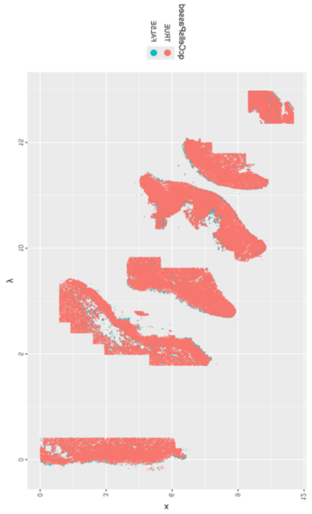

#### Batch Correction

4. **Normalize the expression data and perform dimensionality reduction.**
   4.1. Configure the analysis to run sequentially and increase the allowable memory size for the large CosMx object. Normalize gene-expression values, identify variable genes, scale the data, and calculate the first 30 principal components. These components provide the initial low-dimensional representation used for batch correction.
5. **Correct sample-level batch effects with Harmony.**
   5.1. Apply Harmony using the manually defined sample_slide variable as the batch identifier. Use the first 30 Harmony dimensions to generate a corrected UMAP, construct the nearest-neighbor graph, and identify transcriptional clusters. A clustering resolution of 0.5 is used to generate approximately 20 clusters, although this value may be adjusted depending on the desired level of cell-state resolution.
6. **Assess batch correction and clustering quality.**
   6.1. Preserve the original AtoMx UMAP as a separate reduction and compare the distributions of the six samples before and after Harmony correction. Successful correction should reduce sample-driven separation without eliminating biologically meaningful differences. Visualize the new Seurat clusters in both UMAP and tissue space to identify clusters associated with specific samples, FOV boundaries, or anatomically implausible regions.

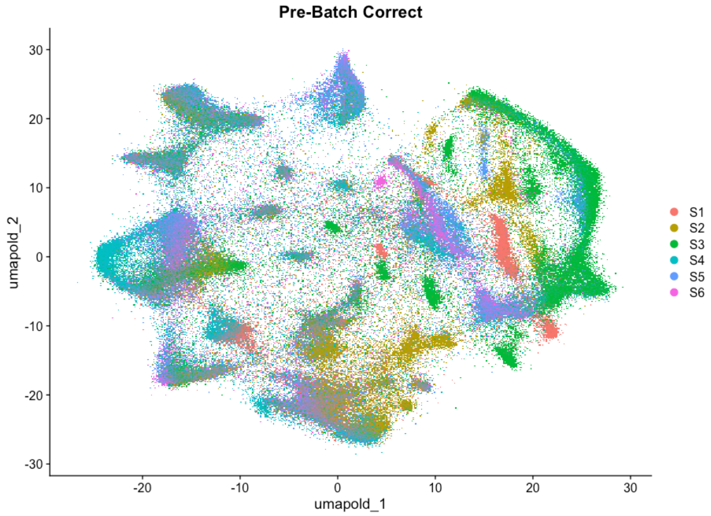

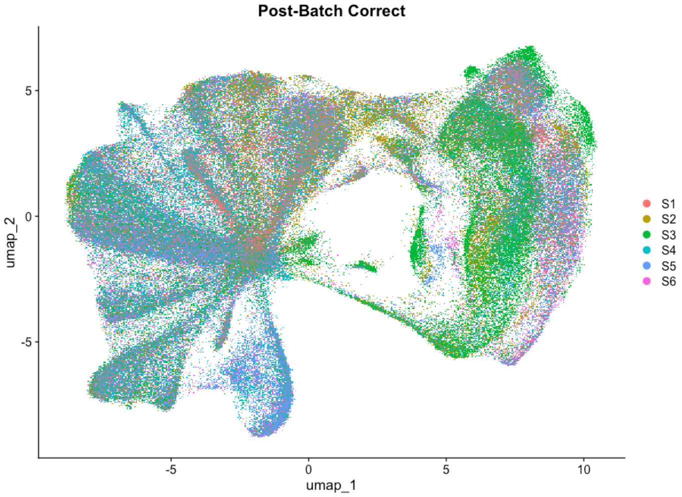

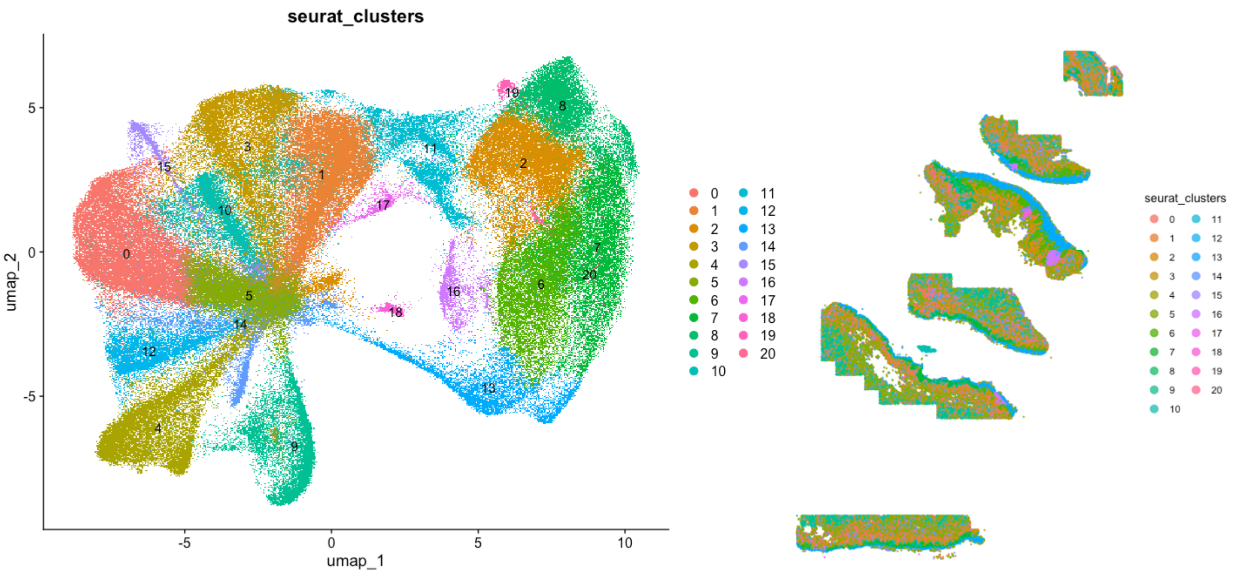

#### Define Cell-types

7. **Identify marker genes for each cluster.**
   7.1. Set the new Seurat clusters as the active identities and use FindAllMarkers() to identify genes positively enriched in each cluster. Retain the top approximately 30 genes ranked by average log-fold change. Review these marker combinations to determine the most likely biological identity of each cluster, using established epithelial, immune, stromal, vascular, and muscle marker patterns.
8. **Assign biological cell-type labels.**
   8.1. Map each numerical cluster to a final cell-type annotation. The principal categories include basal epithelial states, squamous and keratinized epithelial populations, T cells, B cells, macrophages, dendritic cells, mast cells, fibroblasts, endothelial cells, pericytes, and muscle cells. Clusters with mixed or biologically incompatible marker profiles should be labeled conservatively, retained as unknown, or excluded from downstream analyses.

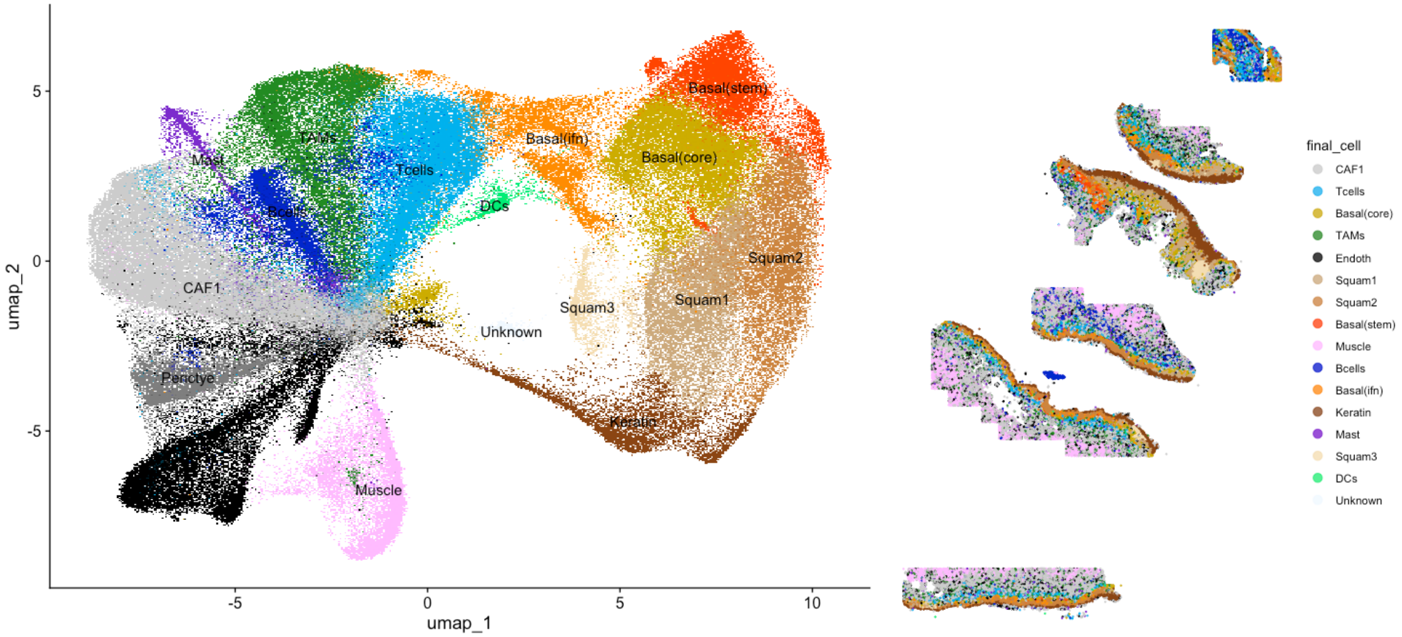

9. **Validate the final annotations.**
   9.1. Visualize the annotated cell types on the Harmony UMAP using a consistent cell-type color palette. Plot the same annotations using the original slide coordinates to confirm that each population occupies a histologically plausible region. Final labels should be supported jointly by marker-gene expression, UMAP structure, and spatial tissue distribution rather than by any single marker.

10. **Save the processed dataset.**
    10.1. Save the final Seurat object after QC filtering, Harmony correction, clustering, and cell-type annotation. This processed object can then be used for downstream spatial analyses, cell-polygon generation, neighborhood analysis, and registration of CosMx data into corresponding H&E image space.

### VIPS: cell polygons and image generation

```
#######################################################################################################################################
#######################################################################################################################################
##     ########      ########      ########     ########              #########           ##       ############      ##
##     ##     ##     ##       ##   ##           ##     ##             ##      ##         ####           ##         ####
##     ##     ##     ##     ##     ##           ##     ##             ##        ##      ##  ##          ##         ##  ##
##     ########      #######       ########     ########              ##          ##   ##    ##         ##        ##    ##
##     ##            ##     ##     ##           ##                    ##        ##    ##########        ##       ##########
##     ##            ##      ##    ##           ##                    ##      ##     ##        ##       ##      ##        ##
##     ##            ##        ##  ########     ##                    #########     ##           ##     ##     ##          ##
#######################################################################################################################################
#######################################################################################################################################
##           ##      ###    ##    #######  ##     ##  #######    #######         ######
##          ####     ####   ##   ##        ##     ## ##     ##   ##    ##        ##
##         ##  ##    ## ##  ##   ##        ##     ## ##     ##   ##    ##        ##
##        ##    ##   ##  ##  ##  ##        ######### ##     ##   #######         ######
##       ##########  ##  ## ##   ##        ##     ## ##     ##   ##    ##             ##
##       ##      ##  ##   ####    ##       ##     ## ##     ##   ##     ##            ##
##       ##      ##  ##    ###    ######## ##     ##  #######    ##      ##      ######
#######################################################################################################################################
#######################################################################################################################################
```

This procedure converts annotated CosMx data into a high-resolution image for H&E registration. A processed Seurat object and cell-polygon file are combined in R to generate a cell-type-colored PNG, which is then converted in the terminal with VIPS into a pyramidal TIFF and exported from QuPath as an OME-TIFF. The final image preserves CosMx spatial geometry and supports anchor selection, tissue-level alignment, and multimodal comparison.

#### Generate High Resolution PNG in R

1. **Load the annotated CosMx object and define the cell-type colors**
   1.1. Begin by loading the previously processed PRESCCT1_HNSC_cellTypes.RData object, which contains the final cell-type annotations in the Seurat metadata.
   1.2. Define a named color vector (simp2color) so that each biological cell type is assigned a consistent fill color. These colors are later used both for plotting the polygons in R and for exporting metadata to the Shiny anchor app.
2. **Load the polygon flat file and prepare the metadata.**
   2.1. Read the CosMx polygon file (PRESCCT2-polygons.csv.gz) using fread(). This file contains the cell boundary coordinates needed to draw the polygon map in CosMx pixel space.
   2.2. Extract the corresponding cell metadata from the Seurat object (nano.obj@meta.data), which contains the final cell-type labels for each cell. At this stage, the metadata are used to confirm the distribution of annotated cell types and to link each polygon to its biological label.
3. **Filter the polygons to cells retained after QC and annotation.**
   3.1. Remove polygons corresponding to cells that were excluded during earlier QC or that do not appear in the final Seurat object. This is done by keeping only polygons whose cell identifier matches a cell_id in the processed metadata. The result is a filtered polygon table containing only high-quality, annotated cells that should appear in the final CosMx image.
4. **Export simplified polygon and metadata tables for the Shiny anchor app.**
   4.1. Create a reduced metadata table containing the cell identifier, FOV, final cell-type label, and assigned plotting color.
   4.2. Save both the filtered polygon table and the simplified metadata table as CSV files. These files are intended for downstream use in the registration workflow, especially in the R Shiny anchor app, where a lightweight polygon representation is preferred for speed and interactive viewing.
5. **Attach cell-type labels directly to the polygon table.**
   5.1. Create a mapping from each cell_id to its final cell-type label and append that label to the polygon table. This allows ggplot2 to fill each polygon directly by cell type.
   5.2. Once the cell-type label has been added to the polygon data frame, the original Seurat object can be removed from memory if needed to reduce RAM usage before plotting the full-resolution image.
6. **Generate the CosMx polygon image in R.**

6.1. Use ggplot2 to plot each cell polygon in global CosMx pixel coordinates (x_global_px, y_global_px), grouping vertices by cell and filling polygons by final_cell. The plot is drawn with geom_polygon(), fixed aspect ratio (coord_fixed()), a blank theme (theme_void()), and a manual fill scale using the previously defined cell-type colors. The legend is removed so that the output is a clean, full-slide cell-type image suitable for registration. This figure represents the CosMx data in its native spatial coordinate system.

6.2. **Save the polygon map as a high-resolution PNG in R.**

- NOTE 1: Uncheck "use device pixel ratio so that output stays as close to native resolution as possible.
- NOTE 2: if exporting from plot window, collapse window-pane to prevent white bars on side of image
- NOTE 3: otherwise no other export precautions are needed as this PNG is justed used as a rough visual reference (to confirm landmark presence in H&E and CosMx). Precise image export into EXACT H&E space will be conducted after the registration step

#### Image Processing in VIPS (white background and tiled format)

7. If necessary flip H&E to match orientation of CosMx when load into QuPath (this is necessary because H&E will be ground truth pixel space)
   - vips flip "PreSCCT1_Series2.ome.tif" \
   - "PreSCCT1_Series2_flipY_pyramid.ome.tif[tile,pyramid,compression=jpeg,Q=90,bigtiff,tile-width=256,tile-height=256]" \
   - horizontal
8. **Install VIPS and run it from the terminal.**

   8.1. If Homebrew is not already installed, install it first from the official Homebrew website and then rerun the brew install vips command. These commands are run in the **Terminal application**, not in the R console.

   8.2. The next steps are performed **outside R**, in the **system terminal**. On macOS, install VIPS with Homebrew using:

   brew install vips
   After installation, confirm it is available by typing:
   vips --version

9. **Use VIPS in the terminal to prepare a pyramidal TIFF.**

   9.1. First, flatten the PNG so that any transparent background becomes solid white. Run the following in the **terminal**:
   vips flatten PRESCCT1_CosMx_raw.png PRESCCT1_CosMx_white.png --background "255 255 255"

   9.2. Next, convert the white-background PNG into a tiled pyramidal TIFF using:
   vips tiffsave PRESCCT1_CosMx_white.png PRESCCT1_CosMx_pyr.tif --tile --tile-width=256 --tile-height=256 --pyramid --compression=lzw
   This creates a pyramidal TIFF that is much easier to open and handle in QuPath than a single massive flat image. At this stage, the file is a pyramidal TIFF but not yet an OME-TIFF.

10. **Open the pyramidal TIFF in QuPath and export it as an OME-TIFF.**\

    10.1. Open the pyramidal TIFF in QuPath, then export it as a full-resolution OME-TIFF.

    10.2. In QuPath, use: **File > Export image as > Original Pixel**

    10.3. Save the image without downsampling so that the full spatial resolution is preserved. This final step converts the VIPS-generated pyramidal TIFF into an OME-TIFF that can be used in downstream histology workflows and registration pipelines.

11. **Final output and purpose.** The overall result of this workflow is a high-resolution cell-polygon image in CosMx space, first saved as a PNG, then converted into a pyramidal TIFF, and finally exported as an OME-TIFF. This image serves as the key input for manual or semi-manual registration workflows, allowing the CosMx-derived cell-type map to be aligned against the corresponding H&E slide in QuPath or related tools.

### QuPath: Defining landmarks conserved across modalities

#### Setup QuPath

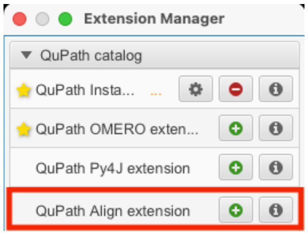

1. Install QuPath: https://qupath.github.io/
2. Open QuPath
3. Install QuPath Align Extension:
   - **3.1. Extensions > Manage Extensions**
   - 3.2. Click + next to QuPath Align Extension

#### Setup Project

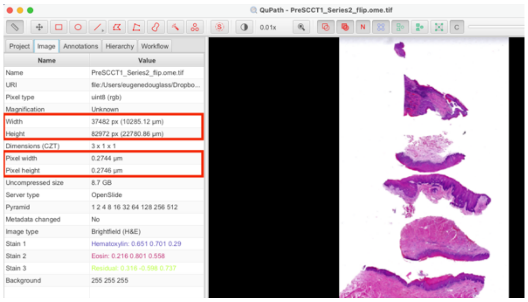

4. Create and Save Project
   - **4.1. File > Project > Create Project**
   - 4.2. Select empty folder: "tmp"
5. Add CosMx and H&E images to Project
   - 5.1. Drag files into Project side bar
   - 5.2. Select Import
6. Roughly match sizes of images
   - 6.1. Double click each image and inspect height and width for each in Image Sidebar Tab and use to estimate approximate length conversion from CosMx -> H&E
     - 6.1.1. **H&E:** 37,483 x 82,972 px
     - 6.1.2. **CosMx:** 28,443 x 44,000 px
     - 6.1.3. **Conversion:** ~2x size
   - 6.2. Record H&E Pixel Width baseline (this value is accurate and CosMx should be adjusted to it)
     - 6.2.1. **H&E:** 0.274 um/px
     - 6.2.2. **CosMx**: 352.9 um/px
   - 6.3. Double click on CosMx pixel width to change values to conversion factor * H&E pixel width
     - 6.3.1. 2*0.274 = 0.548 um/px ~ CosMx (rough)

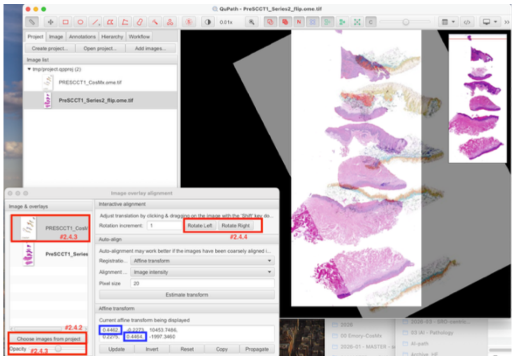

7. Fine tune size-match with Align Extension
   - 7.1. Set H&E as baseline image by double clicking on its icon in the Project tab of the side-bar
   - 7.2. Open QuPath Align Extension:
     - 7.2.1. **Analyze > Alignment > Interactive Image Alignment**
     - 7.2.2. Click "Choose images from Project"
     - 7.2.3. Move CosMx image from "Available" to "Selected"
   - 7.3. Click on top layer "CosMx" and set Opacity to ~50%
   - 7.4. Click rotate buttons until layer-orientation is matched
   - 7.5. Iteratively adjust placement and scale
     - 7.5.1. **Move overlap using**: Hold Shift + Click on Viewer and drag to overlay one section
     - 7.5.2. **Adjust scale using**: Affine **diagonal elements (in blue)** to scale up and down (both elements should be exactly the same as they reflect vertical and horizontal scaling)
     - 7.5.3. **Rotate using:** rotate buttons as needed
     - 7.5.4. **Repeat** until one section is "perfectly" overlaid across both layers

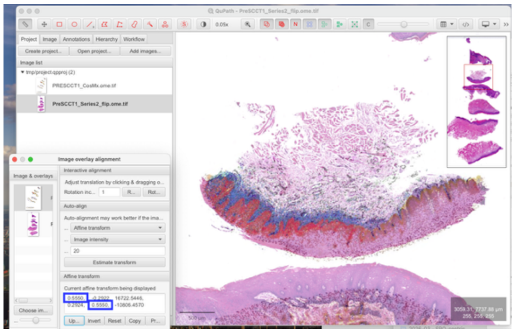

   - 7.6. Zoom out and readjust rotation ONLY back to approximately starting point. Record scale from the affine diagonal, this is your final CosMx/H&E Conversion Factor
     - 7.6.1. Affine diagonal scale: 0.627
   - 7.7. Convert previous CosMx um/px to new value using conversion factor above
     - 7.7.1. 0.278 / 0.627 = **0.437 um/px** = CoxMx
   - 7.8. **Close Alignment** window
   - 7.9. **Save Project** (CosMx and H&E scales are roughly matched so you will just have to rotate and translate in future allignments)

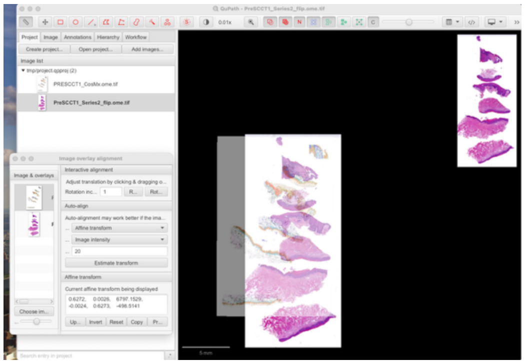

8. Add Pathologist Annotations
    - 8.1. **Drag Pathologist annotation** on top of H&E (while open in main window) and **Save**

#### Setup Workspace:

9. Open Project
10. Align One Section H&E and CosMx
    - 10.1. Set H&E as baseline image by double clicking on its icon in the Project tab of the side-bar
    - 10.2. Open QuPath Align Extension:
        - 10.2.1. **Analyze > Alignment > Interactive Image Alignment**
        - 10.2.2. Click “Choose images from Project”
        - 10.2.3. Move CosMx image from “Available” to “Selected”
    - 10.3. Click on top layer “CosMx” and set Opacity to ~50%
    - 10.4. Iteratively adjust placement and rotation (scale should be fine based on Step #2)
        - 10.4.1. **Rotate using:** rotate buttons as needed
        - 10.4.2. **Repeat** until one section is “perfectly” overlaid across both layers
11. Close Side Panel
    - **11.1. View > Show Analysis Pane (uncheck)**
12. Shrink Image Overlay Alignment Window and Move to Left
13. Add Points Window Panel to left side
14. Add 10 points labeled L1-L10 within points panel

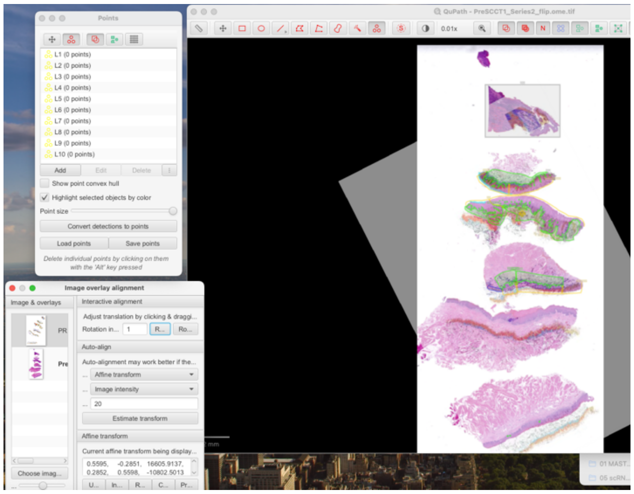

#### Define/Save Anchors per sample

15. Zoom into section of interest
16. Set Transparency to 50%
17. **Rotate/Rescale for each section**
18. Coarse-Allign Right Hand side (Shift + drag) to tissue border and 1-2 internal features with:
        - 18.1.1. **preferably annotated features**
        - 18.1.2. **preferably macroscopic geometry (circular > angled > flat)**
19. Zoom into Right Hand Side & set landmarks
    - 19.1. Fine-Align Right hand side (shift drag) to tissue border and 1-2 internal features
    - 19.2. Set 1-2 macroscopic anchors
    - 19.3. Toggle between full H&E and CosMx to confirm Anchors on visible landmarks in both modalities

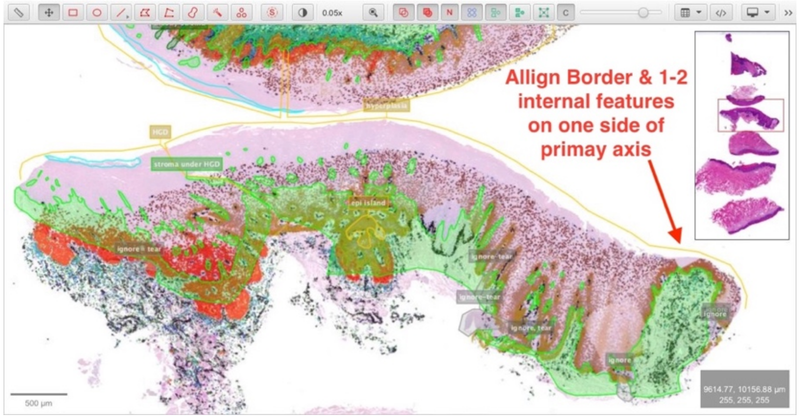

20. Zoom in Further on Finer grained Landmarks
    - 20.1. ducts, vessels, islands (BEST)
        - 20.1.1. **Fidelity:** unambiguous geometry match (esp when combined with tissue borders)
        - 20.1.2. **Anchor:** center is easy anchor (less variable than borders)
    - 20.2. Corners, Tips (2nd BEST)
        - 20.2.1. **Fidelity:** clear geometry match (especially with pathologist annotation)
        - 20.2.2. **Anchor:** ok, sometimes difficult if not point

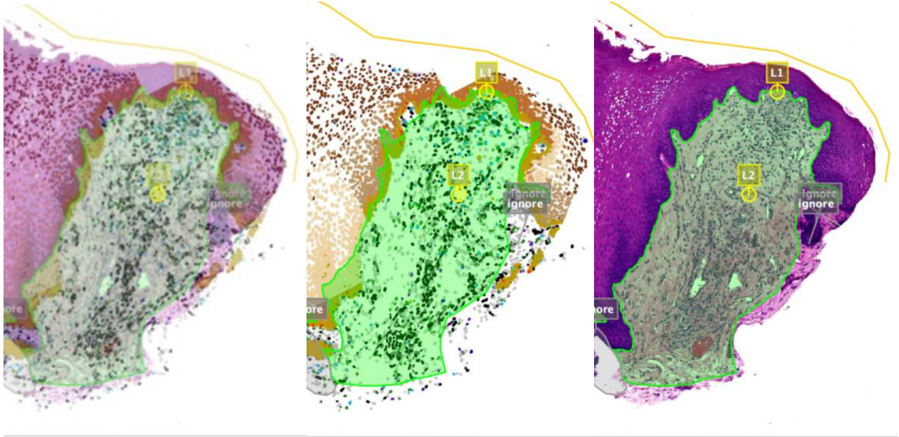

21. Repeat for Left Side
22. Repeat for center

22.1.1. **NOTE:** for ambiguous landmarks toggle between right- and left- alignment, circular objects like vessles in CosMx layer will “circle” the H&E feature (as left and right anchors pull to one side or the other)

23. SAVE ALL POINTS AS TSV FILE
24. TAKE SCREENSHOTS AT EVERY STAGE (100% CosMx and 100%HE w/ Anchor + annotations):

### QuPath: Defining Tissue Borders in H&E slides

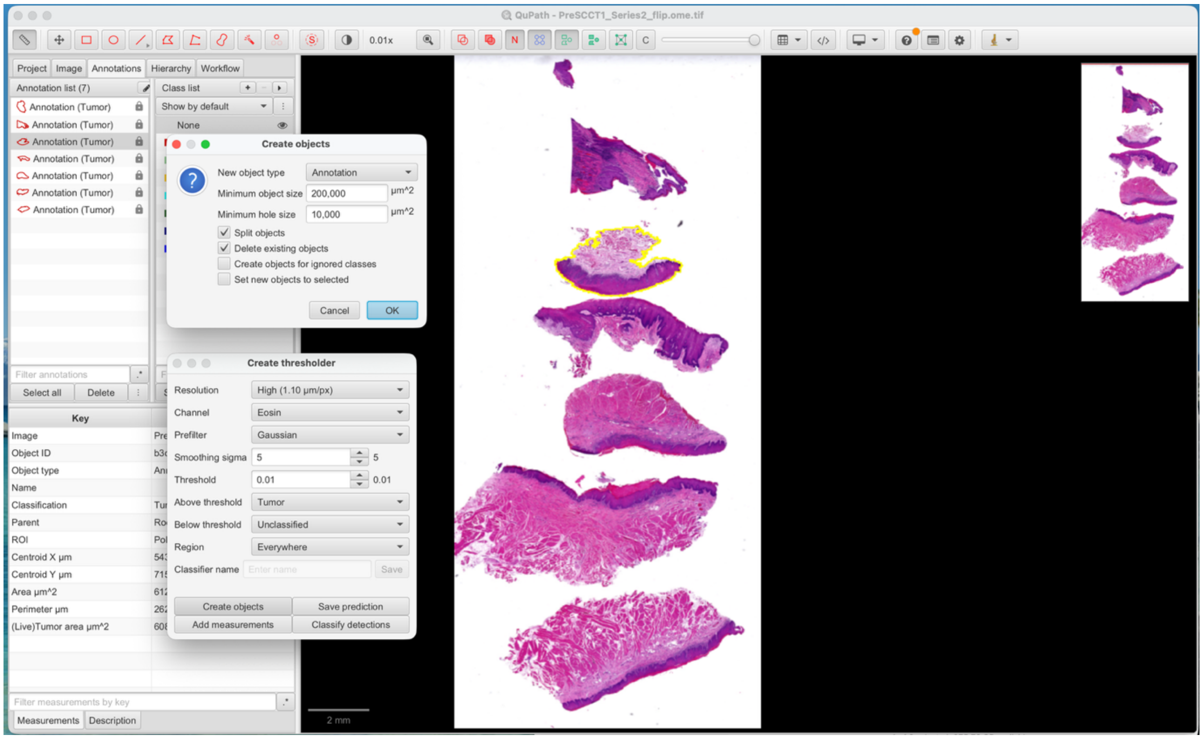

1. Open the H&E OME-TIFF in QuPath and select **Classify** → **Pixel classification** → **Create thresholder**. Thresholding is used to separate stained tissue from the white background and convert the detected regions into annotations.

2. Configure the thresholder using the settings shown in the screenshot: **Resolution: High (1.10 µm/px)**, **Channel: Eosin**, **Prefilter: Gaussian**, **Smoothing sigma: 5**, and **Threshold: 0.01**. Assign pixels **above the threshold** to **Tumor**, pixels **below the threshold** to **Unclassified**, and set **Region: Everywhere**.

3. Inspect the colored preview overlay and confirm that it follows the outer tissue borders while excluding the white slide background. Adjust the threshold only when necessary for a different slide.

4. Select **Create objects** and apply the classifier across the full image. In the object-creation window, use **New object type: Annotation**, **Minimum object size: 200,000 µm²**, and **Minimum hole size: 10,000 µm²**.

5. Enable **Split objects** so spatially separate tissue sections become individual annotations. Enable **Delete existing objects** when replacing a previous border set. Leave **Create objects for ignored classes** and **Set new objects to selected** unchecked.

6. Click **OK**, visually inspect the resulting tissue-border annotations, and manually correct any missed tissue, debris, or inappropriate holes before exporting the annotations for registration.

### AnchorR: defining CosMx landmarks and Affine Fit

#### Setup Shiny App

1. **Install the required R packages**
   1.1. Open R or RStudio and install the packages required by AnchorR:
   1.2. install.packages(c("shiny", "ggplot2", "data.table", "scales", "png", "sf", "dplyr", "ragg"))
   1.3. Place the app.R file in a dedicated folder. AnchorR runs locally as an R Shiny application and does not require specialized computer-vision software.
2. **Prepare the CosMx input files**
   2.1. Place cell_polygons.csv and cell_meta.csv (Outputs from VIPS step) in the same folder as app.R. The polygon file must contain the columns cell, fov, x_global_px, and y_global_px. The metadata file must contain cell and fov. Including final_cell and color allows the application to display the intended cell-type labels and colors. An optional cell_type_key.jpg can be placed in the application folder or in a www subfolder.

#### Set CosMx Anchors (Tab #1):

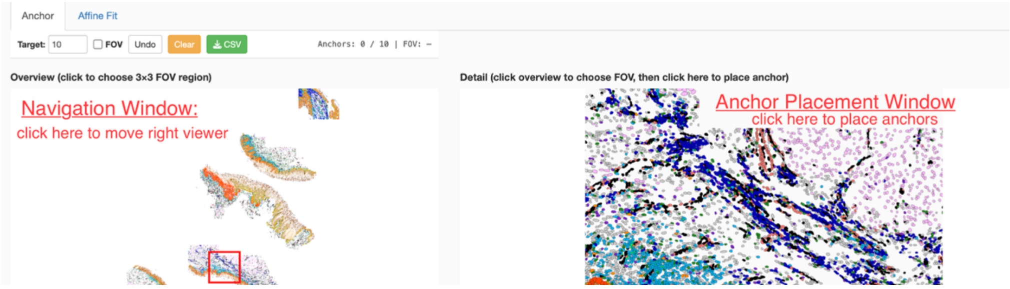

3. **Launch AnchorR**
   3.1. Open app.R in RStudio and click Run App. Alternatively, set the working directory to the application folder and run:
   3.2. shiny::runApp()
   3.3. The application automatically loads the CosMx input files and generates a cached whole-slide overview. The initial rendering may take approximately 30–60 seconds or longer for a large dataset, but subsequent sessions reuse the cached image.
4. **Setup QuPath vs Anchor R work-space (screenshot below)** that juxtaposes screenshots from QuPATH with CosMx overlay with app. This helps user rapidly click through 10 anchors per section

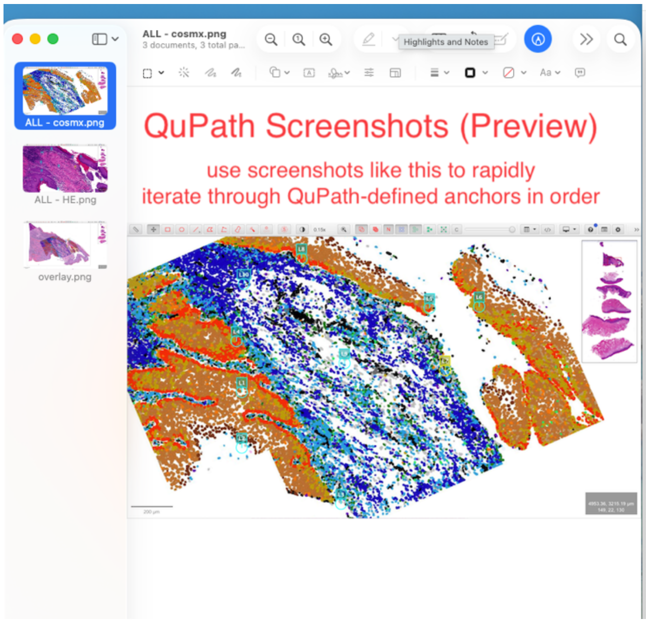

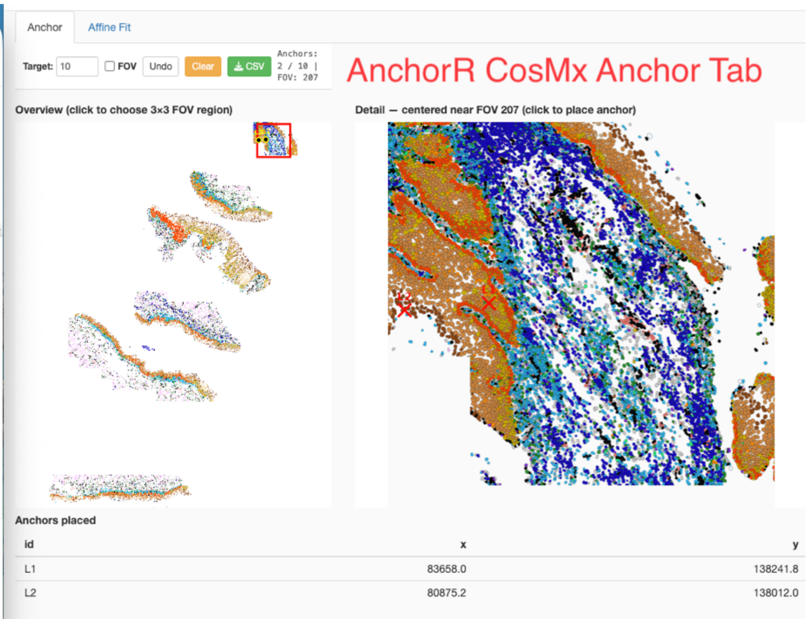

5. **Navigate the CosMx map in the Anchor tab**
    5.1. Set the target number of anchors, typically 10 per tissue section.
    5.2. Click a location in the Overview panel to center the detailed view on that region. The Detail panel displays the selected area and the surrounding 3 × 3 field-of-view neighborhood.
    5.3. Select the FOV option to display field boundaries and identifiers.
6. **Place CosMx landmarks**
    6.1. Click the precise landmark location in the Detail panel. Anchors are labeled sequentially as L1, L2, and so forth.
    6.2. Distribute anchors across the full tissue section, including both ends and the center.
    6.3. Prioritize distinctive internal structures such as vessels, ducts, epithelial islands, and epithelial-stromal boundaries.
    6.4. Use corners or tissue tips only when stronger internal landmarks are unavailable. Each landmark should be confirmed as visible in both the CosMx and H&E images.
7. **Review and export the CosMx anchors**
    7.1. Use Undo to remove the most recently placed anchor or Clear to remove all anchors. Review the anchor table, then select CSV to export the coordinates. The resulting file contains the columns id, x, and y. The anchor identifiers must match the corresponding H&E landmark names exported from QuPath.

#### Calculate and Refine Affine Fit

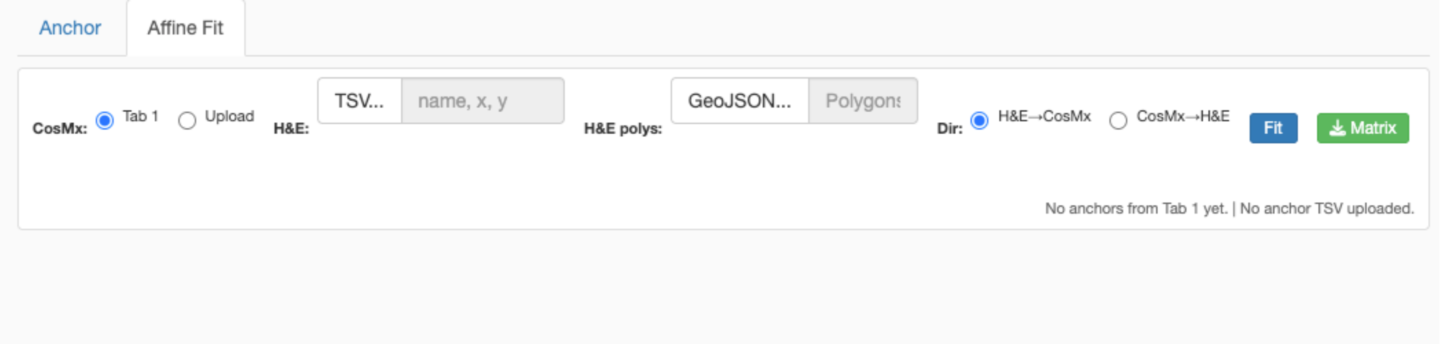

8. **Load the registration inputs in the Affine Fit tab**
    8.1. Use the anchors directly from the Anchor tab or upload a previously exported CosMx anchor CSV. Upload the H&E landmark file exported from QuPath as a TSV, TXT, or CSV file containing the columns name, x, and y. The name values must match the CosMx anchor identifiers. Optionally upload the H&E tissue borders as a GeoJSON file for visual quality control. At least three paired anchors are required, although approximately 10 well-distributed anchors are recommended.
9. **Fit the affine transformation**
    9.1. Select the transformation direction and click Fit. Use CosMx to H&E when the goal is to transform CosMx cell polygons into H&E coordinates. Use H&E to CosMx when the inverse mapping is required. AnchorR matches landmarks by identifier and estimates a two-dimensional affine transformation using least-squares fitting.
10. **Evaluate and refine the registration**
    **10.1.** Review the reported root-mean-square error, maximum residual, number of included anchors, determinant, translation, rotation, scale, and shear. Inspect the predicted-versus-observed landmark plot, residual scatter plot, and per-anchor residual table. Uncheck a poorly matched anchor to exclude it and refit the model. When an H&E border GeoJSON is supplied, inspect the transformed tissue outline over the CosMx cell-type map to confirm visually plausible tissue-level alignment.

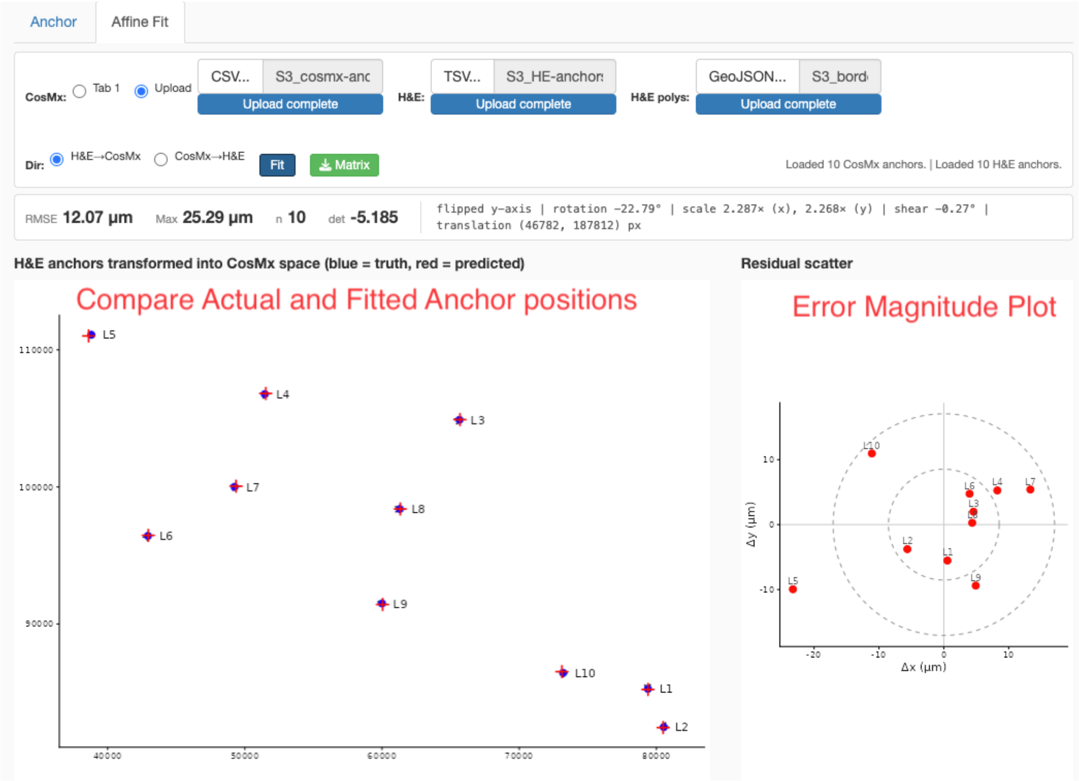

**11. Export the final registration outputs**

12. Select Matrix to export the fitted 3 × 2 affine transformation matrix. The application can also export transformed H&E borders as GeoJSON and save the registration overlay as a PNG. Repeat the complete fitting procedure independently for each tissue section because section-specific affine transformations better capture differences in orientation, scale, translation, and shear than a single whole-slide transformation.

### Finalize CosMx polygons and images in H&E space

#### Load Data

1. Load the annotated CosMx data and polygon coordinates
    1.1. Load the processed Seurat object containing final cell-type and tissue-section annotations. Import the original CosMx polygon flat file using fread(). The polygon file provides the boundary vertices for each cell in global CosMx pixel coordinates, while the Seurat metadata provides the corresponding cell type and sample identity.
2. Filter the polygon file
    2.1. Retain only polygons whose cell identifiers are present in the quality-controlled Seurat object. This removes polygons belonging to cells that failed QC or were excluded during cell-type annotation.
3. Add cell-type labels and colors
    3.1. Create a mapping between each cell identifier and its final cell-type annotation, then add the resulting final_cell column to the polygon table. Use the existing simp2color palette to maintain consistent cell-type colors across the CosMx and H&E-space images.
4. Verify the polygons in native CosMx space
    4.1. Plot the polygons using x_global_px and y_global_px, grouping vertices by cell and filling each polygon by final_cell. Use a fixed aspect ratio and remove axes and legends. This plot confirms that the filtered polygon file and cell-type annotations were joined correctly before registration.

#### Transform CosMx Polygons into H&E Space

5. Assign each polygon to a tissue section
    5.1. Transfer the sample_slide annotation from the Seurat metadata to the polygon table and store it as sample. Plot the polygons colored by sample to confirm that cells have been assigned correctly to sections S1–S6. Each section will receive its own affine transformation.
6. Apply the section-specific affine transformations
    6.1. Create empty x_he_px and y_he_px columns for the transformed coordinates. For each tissue section, load the corresponding CosMx-to-H&E affine matrix, such as S1_cos2he.csv. Extract all polygon vertices belonging to that section and arrange their coordinates as:
    6.2. X = [x_global_px, y_global_px, 1]
    6.3. Multiply these coordinates by the 3 × 2 affine matrix:
    6.4. Predicted H&E coordinates = X × A
    6.5. Store the resulting coordinates in x_he_px and y_he_px. Repeat this process independently for all six tissue sections.
7. Inspect the transformed polygons
    7.1. Plot the transformed polygons using x_he_px and y_he_px. Reverse the y-axis because H&E images use a top-left origin, whereas standard R plots use a bottom-left origin. Confirm that all sections have plausible positions, orientations, scales, and spatial relationships before exporting the final image.
8. Save the transformed polygon dataset
    8.1. Save the final cell_polygons object as cell_polygons.RData. This file retains the original CosMx coordinates, H&E-space coordinates, cell identifiers, tissue-section labels, and final cell-type annotations. It can be used for downstream spatial analyses directly in H&E coordinate space.

#### Export CosMx Colored Cells as Image (in H&E pixels)

9. Match the output canvas to the H&E image
    9.1. Set the plotting limits to the exact pixel dimensions of the H&E whole-slide image. In this example:
    9.2. W = 37,482 pixels
    H = 82,972 pixels
    9.3. Set the x-axis limits to 0–W and reverse the y-axis from H to 0. Remove all plot margins and preserve a fixed aspect ratio. This ensures that the exported CosMx image occupies the same coordinate system and canvas dimensions as the H&E image.
10. Export the registered CosMx image as a PNG

10.1. Full-resolution rendering may exceed available memory. If necessary, downsample the output image while retaining the original H&E coordinate limits. For a threefold downsampling:

10.2. s = 3
Output width = 12,494 pixels
Output height = 27,657 pixels

10.3. Export the image using ragg::agg_png with a white background and limitsize = FALSE. Although the raster image is downsampled, the plot continues to represent the full H&E coordinate range.

11. Convert the PNG to a pyramidal TIFF using VIPS

11.1. Run VIPS from the system Terminal, not from the R console. Convert the PNG into a tiled pyramidal TIFF using:

11.2. vips tiffsave PRESCCT1_CosMx_HEreg_raw.png PRESCCT1_CosMx_HEreg.tif
--tile --tile-width=256 --tile-height=256 --pyramid
--compression=lzw

11.3. The PNG already has a white background, so a separate flattening step is not required.

12. Convert the TIFF to OME-TIFF in QuPath

12.1. Open the pyramidal TIFF in QuPath and export it at full available resolution as an OME-TIFF using:

12.2. File > Export image as > Original Pixels

12.3. The resulting image can be directly overlaid with the H&E image because both use the same coordinate system and spatial orientation.

13.

### OpenSeaDragon Webinterface

#### Prepare Files

1. Prepare the registered images

   - Begin with the final H&E and CosMx images in the same coordinate system, orientation, and canvas dimensions.
   - Use pyramidal OME-TIFF files when available.
   - Assign simple filenames, such as:
     - PRESCCT1_HE.ome.tif
     - PRESCCT1_CosMx.ome.tif
     - PRESCCT1_IF.ome.tif

2. Install VIPS

   - Run the following commands in the macOS Terminal, not in R or RStudio.
   - Install VIPS using Homebrew: brew install vips
   - Confirm that VIPS was installed correctly: vips --version

3. Convert each image to Deep Zoom Image format

   - In the Terminal, navigate to the folder containing the images: cd /path/to/viewer-folder
   - Convert each image using vips dzsave:
     - vips dzsave PRESCCT1_HE.ome.tif HE
     - vips dzsave PRESCCT1_CosMx.ome.tif CosMx
     - vips dzsave PRESCCT1_IF.ome.tif IF
   - Each command generates:
     - A DZI descriptor file, such as HE.dzi
     - A corresponding tile folder, such as HE_files
   - Keep each .dzi file in the same directory as its associated tile folder.

4. Prepare the OpenSeadragon application folder

   - Create a folder containing:
     - The .dzi descriptor files
     - The corresponding tile folders
     - The HTML viewer file
     - Any legends, GeoJSON annotations, or interface assets
   - Load OpenSeadragon either from a public content-delivery network or from a local copy of openseadragon.min.js.
   - When using a local copy, also include the OpenSeadragon images folder containing the navigation icons.

#### Build/Modify HTML Viewer

5. Create the HTML viewer

   - Create an HTML file named annotations.html or viewer.html.
   - Define two side-by-side viewer panels, with one showing the H&E image and the other showing the CosMx image.
   - Initialize the viewers using the corresponding DZI files
   - Example code:

```
var viewer1 = OpenSeadragon({
id: "viewer1",
prefixUrl: "https://openseadragon.github.io/openseadragon/images/",
```

```
tileSources: "HE.dzi"
});

var viewer2 = OpenSeadragon({
id: "viewer2",
prefixUrl: "https://openseadragon.github.io/openseadragon/images/",
tileSources: "CosMx.dzi"
});
```

- Set the panel dimensions using CSS. For example, each viewer can occupy approximately 48% of the page width with a height of 800 pixels.

6. Synchronize the two viewers

- Add JavaScript handlers that copy the zoom level and viewport center between the viewers.
- Synchronize both directions so that either viewer can control the other:

```
function syncViewers(viewerA, viewerB) {
viewerA.addHandler("animation", function() {
viewerB.viewport.zoomTo(
viewerA.viewport.getZoom(),
null,
true
);

    viewerB.viewport.panTo(

      viewerA.viewport.getCenter(),

      true

    );

  });

}

syncViewers(viewer1, viewer2);

syncViewers(viewer2, viewer1);
```

- Confirm that the images remain aligned while zooming and panning.

#### Test Locally

7. Test the application locally

- Browsers generally cannot load DZI tiles when the HTML file is opened directly from the filesystem.
- In the Terminal, navigate to the application folder: cd /path/to/viewer-folder
- Start a local web server: python3 -m http.server 8000
- Open the application in a browser at: http://localhost:8000/annotations.html
- Confirm that:
    - Both images load correctly
    - Zooming and panning are synchronized
    - Registered tissue structures remain aligned
    - Legends and annotations display correctly

8. Configure remote storage for online deployment

- Store the large DZI tile folders in object storage rather than on a conventional website host.
- Cloudflare R2 can be used for this purpose.
- Install rclone in the Terminal: brew install rclone
- Configure the Cloudflare R2 connection: rclone config
- During configuration:
- Create a new remote
- Select s3 as the storage type
- Select Cloudflare as the provider
- Enter newly generated R2 access credentials
- Enter the R2 endpoint associated with the account
- Do not include access keys or secret keys in the HTML, source code, or shared documentation.
- Test the connection:  rclone lsd r2:your-bucket

#### Deploy Online using CloudFlare

9. Upload the DZI files and tiles

- Upload the complete project folder using:

```
rclone sync "/path/to/local/project-folder" r2:your-bucket/project-name --transfers 32 --progress
```

- Confirm that the upload includes:
    - Each .dzi file
    - Every associated tile folder
- If some files fail because of temporary server errors, rerun the same command. Completed files will be skipped and only missing files will be retried.

10. Enable public access and CORS

- Enable a public development URL or custom public domain for the Cloudflare R2 bucket.
- Add a Cross-Origin Resource Sharing policy that allows GET requests from the website hosting the OpenSeadragon interface.
- Restrict the allowed origins to the intended website domains. For example:

```
[
  {
    "AllowedOrigins": [
      "https://example.com",
      "https://www.example.com"
    ],
    "AllowedMethods": ["GET"],
    "AllowedHeaders": ["*"]
  }
]
```

- Without the correct CORS policy, the HTML page may load while the browser blocks the DZI files or image tiles.

11. Update the HTML to use remote image URLs

- Replace relative DZI paths such as: tileSources: "HE.dzi"
- With the complete public URL: tileSources: "https://public-storage-domain/project-name/HE.dzi"
- Update every DZI reference in the application, including:
  - Viewer tileSources
  - Image-selection controls
  - Initial image assignments
  - Legend-selection functions
- When identifying an image from a complete URL, use partial filename matching rather than exact string equality:

```
var isHE = dziValue.indexOf("HE.dzi") !== -1;
```

- OpenSeadragon can be loaded from a CDN using:

```
<script src="https://cdnjs.cloudflare.com/ajax/libs/openseadragon/4.1.0/openseadragon.min.js"></script>
```

12. Upload the web interface

- Upload the lightweight application files to the main website host:
  - annotations.html
  - JavaScript and CSS files
  - Cell-type legends
  - GeoJSON annotations
- OpeSeadragon navigation icons, when not using a CDN
- Keep the large DZI tiles in object storage.
- Upload the website files to the web-accessible directory, such as:
- public_html/project-name/
- Open the final public URL and test the interface on both desktop and mobile browsers.
